# Mitochondrial state determines functionally divergent stem cell population in planaria

**DOI:** 10.1101/2020.07.29.226365

**Authors:** Mohamed Mohamed Haroon, Vairavan Lakshmanan, Souradeep R. Sarkar, Kai Lei, Praveen Kumar Vemula, Dasaradhi Palakodeti

## Abstract

Mitochondrial state changes were shown to be critical for stem cell function. However, variation in the mitochondrial content in stem cells and the implication, if any, on differentiation is poorly understood. Here, using cellular and molecular studies, we show that the planarian pluripotent stem cells (PSCs) have low mitochondrial mass compared to its progenitors. Further, the mitochondrial mass correlated with OxPhos and inhibiting the transition to OxPhos dependent metabolism in cultured cells resulted in higher PIWI-1^High^ neoblasts. Transplantation experiments provided functional validation that neoblasts with low mitochondrial mass are the true PSCs. In summary, we show that low mitochondrial mass is a hallmark of PSCs in planaria and provide a mechanism to isolate live, functionally active, PSCs from different cell cycle stages (G0/G1 and S, G2/M). Our study demonstrates that the change in mitochondrial metabolism, a feature of PSCs is conserved in planaria and highlights its role in organismal regeneration.

## Introduction

Neoblasts are adult pluripotent stem cells that play a central role in planarian regeneration. They are capable of differentiating into all other cell types thereby facilitating regeneration of lost tissues. Upon injury, planarians induce wound healing response leading to stem cell proliferation and differentiation, followed by remodeling of old and new tissue to form a completely regenerated animal (Reddien, 2018). In planarians, neoblasts are the sole proliferating cells and eliminating them by irradiation results in lethality (Newmark and Sánchez Alvarado, 2000; Reddien and Alvarado, 2004). Our understanding of the mechanisms that regulate pluripotency is limited by the capacity to isolate the pluripotent stem cells (PSCs) from planaria.

The current technique to isolate neoblasts involve fluorescence-activated cell sorting (FACS) based on the nuclear density by staining the cells with Hoechst (Hayashi et al., 2006). Hoechst staining identified three major populations - X1, X2 and Xins. X1 cells are the proliferating neoblasts with >2N nuclear content (S and G2/M phases of cell cycle) and expression of stem cell markers such as *piwi-1, vasa-1, bruli* (Reddien et al., 2005; Wagner et al., 2012). X2 includes lineage specified progenitor cells as well as stem cells in the G0/G1 phase of cell cycle (Eisenhoffer et al., 2008; Molinaro and Pearson, 2016; Van Wolfswinkel et al., 2014). Both X1, X2 are eliminated upon X-ray irradiation, whereas, the Xins population is insensitive to irradiation and constitutes terminally differentiated cells. Since the neoblast are the only dividing cells in planaria, X1 population, has been extensively used for functional characterization of the neoblast. So far it has not been possible to isolate stem cells from the X2 gate. Therefore, the current knowledge about the neoblasts was obtained by studying X1 population (>2N cells).

Detailed molecular analysis showed that X1 population consist of pluripotent (clonogenic neoblasts) and primed stem cells (specialized neoblasts) (Fincher et al., 2018; Plass et al., 2018; Wagner et al., 2011; Van Wolfswinkel et al., 2014). Clonogenic neoblasts were shown to rescue planarians upon lethal irradiation. Recently, it was shown that a subset of neoblast, which express tetraspanin-1 (TSPAN-1) are pluripotent in nature. Single cell transplantation of TSPAN-1^+^ cells in lethally irradiated planarians rescued the host with greater efficiency validating its pluripotency (Zeng et al., 2018). The specialized neoblasts are defined by the expression of transcripts pertaining to specific lineages in addition to pan neoblast markers such as *piwi-1*. However, lack of strong evidence to show that all the pluripotent stem cells are TSPAN^+^ suggests that there could be a possibility of subset of pluripotent stem cells that lack the expression of TSPAN.

Mitochondria, a hub for cellular energetics has been shown to be critical for stem cell maintenance and differentiation (Folmes et al., 2012; Xu et al., 2013). Evidences show that mitochondria in PSCs exists in a discontinuous fission state with very low oxidative phosphorylation (OxPhos). Several studies have used differences in the mitochondrial activity as a means to demarcate stem cells from their respective progenitors. For example, hematopoietic stem cells (HSC) with low MitoTracker™ Green (MTG) staining exhibited enhanced stemness and showed greater reconstitution potential compared to its MTG High counterpart (Romero-Moya et al., 2013). Similarly, tetramethylrhodamine methyl ester (TMRM) a mitochondrial membrane potential sensing dye was used to separate minimally differentiated stem cell-memory T cells from highly differentiated effector memory T cells (Sukumar et al., 2016).

This study aims to understand the differences in mitochondrial content between the planarian cell populations, its functional consequence on stemness and employ this as a means to distinguish stem cell states. Based on the mitochondrial dye MTG, we were able to further distinguish cells within the X1 and the X2 cell populations as X1-MTG^High^, X1-MTG^Low^, X2-MTG^High^ and X2-MTG^Low^ cells. Transcriptome analysis and molecular characterization of these cells revealed that the X1 cells with low mitochondrial mass were more homogenous pool of pluripotent stem cells compared to cells with higher mitochondrial mass. The X1-MTG^High^ cells expressed genes essential for lineage commitment exhibiting the signatures of committed neoblasts. Similar analysis of X2 revealed that the cells with low mitochondrial mass were either stem cells in G0/G1 phase or early progenitors, whereas the cells with high mitochondrial mass were late progenitors. Our study also revealed that the mitochondrial activity correlates with mass and inhibiting the mitochondrial activity by treating the neoblast with FCCP resulted in higher expression of PIWI-1 *in vitro*, suggestive of increased stemness. Further, to validate the pluripotency, cells isolated from the asexual strain (donor) were transplanted into lethally irradiated sexual strain (host). Our results indicate that the X1 and a subset of X2 cells with low MTG were able to rescue and thus transform the sexual planaria into an asexual one at higher efficiencies compared to the MTG^High^ cells. Together, our results show that the pluripotent stem cells in the planarians can be categorized by low mitochondrial mass compared to their immediate progenitors. Further, the changes in the mitochondrial mass between pluripotent neoblast and the committed neoblast could be used to isolate pluripotent stem cells from planarians for functional characterization.

## Results

### Mitochondrial staining reveals distinct mitochondrial mass in planarian X1, X2, Xins cells

Single cell transcriptome studies in planarians revealed neoblasts are heterogenous pool of clonogenic and specialized stem cells. Here, we investigated how the mitochondrial mass changes between different planarian cell populations. This was tested by staining the neoblast population with MTG, a fluorescent dye that is used as a measure for the mitochondrial mass (Doherty and Perl, 2017; Romero-Moya et al., 2013). The cell suspension of planaria was stained with Hoechst 33342 (nuclear dye) and MTG. The cells stained with Hoechst showed three distinct population, X1, X2 and Xins based on their nuclear content (Figure 1A). X1 cells, which are the proliferating stem cells have lower MTG signal than the Xins population (differentiated cells) (Figures 1B and C). The X2 population, majorly neoblast progenitor had broad distribution of the MTG signal (Figure 1B). MTG fluorescence in X1 cells was more perinuclear and exhibited polarized distribution in the cytoplasm (Figure 1D). Conversely, MTG fluorescence in Xins cells was evenly distributed in the cytoplasm (Figure 1D). To test whether the low MTG intensity in X1 and X2 cells was a consequence of dye efflux, the cells were treated with Verapamil, an efflux pump inhibitor and stained with MTG. There was no significant increase in the median fluorescence intensity in X1 population upon treatment with Verapamil (Figure 1E) suggesting that the lower MTG fluorescence in X1 cells is a true indicator of decreased mitochondrial mass. Together, these results suggest that the difference in the mitochondrial mass between X1 and Xins cells could be an indicator of the stem vs differentiated state.

**Figure 1.**
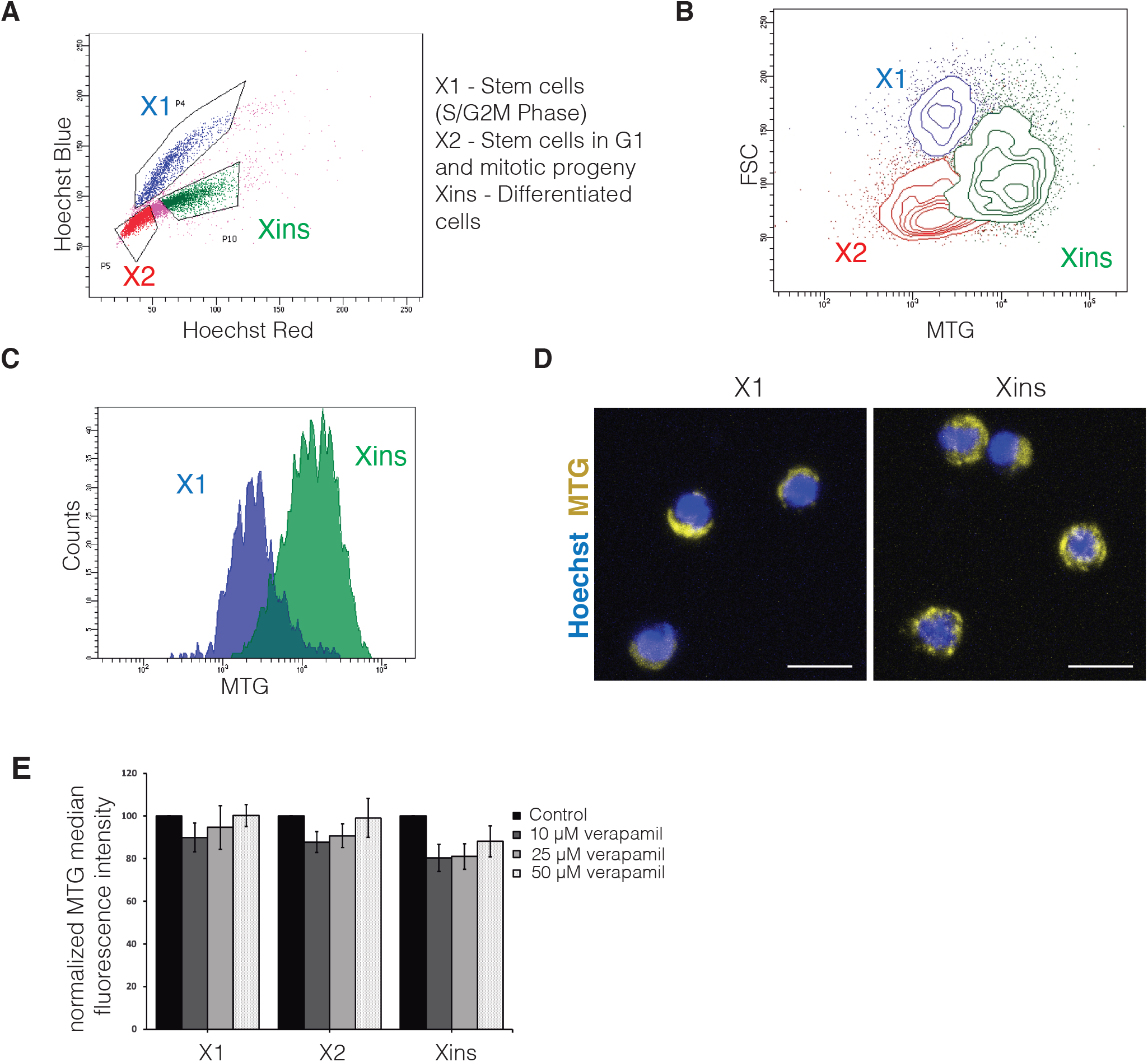
MitoTracker™ Green FM staining in planarian cells. **(A)** Flow cytometry analysis of planarian cell suspension stained with Hoechst 33342 revealing X1, X2 and Xins population. **(B)** and **(C)** Representative contour plot **(B)** and histogram **(C)** showing MTG intensity of X1, X2 and Xins cells. **(D)** Representative confocal images showing MTG staining in sorted, live X1 and Xins cells. Scale bar 10 μm. **(E)** Flow cytometry analysis of MTG median fluorescent intensity of X1, X2 and Xins cells in the presence of indicated amounts of Verapamil. N=3, error bar represents s.e.m.

It has been shown that the mammalian embryonic stem cells show distinct mitochondrial morphology and its activity compared to their progenitors (Xu et al., 2013). However, the changes in the mitochondrial content and its regulation of pluripotency was poorly understood. We examined whether mitochondrial mass could distinguish the clonogenic and specialized neoblast pools. To this end, molecular and functional studies were performed in X1 cells with high and low mitochondrial mass as measured by MTG.

### Low MTG enriches for PIWI-1^High^ cells within X1 population

In addition to the change in the mitochondrial mass between X1 and Xins, we also identified cells with high and low MTG signal within the X1 population (Figure 2A). Here, we wanted to investigate the pluripotent state of the MTG^Low^ and MTG^High^ cells in X1 population. *Piwi-1* is a nuage related gene which has been extensively used to mark stem cells in planaria. To examine the status of *piwi-1* in MTG^Low^ and MTG^High^ cells, we first generated and characterized an antibody to PIWI-1 (Guo et al., 2006). Previous studies have shown that clonogenic neoblast and a subset of neoblast population have high levels of PIWI-1 protein and its corresponding RNA, whereas the progenitors have low levels of PIWI-1 protein (Zeng et al., 2018). The immuno-staining with PIWI-1 antibody in planarian cells identified High, Low and negative PIWI-1 cells (Figure S1A and B). Further, we have also shown that PIWI-1^High^ cells are the proliferating cells, which are in S, G2/M phase whereas the majority of the PIWI-1^Low/Negative^ cells are in G1 phase as was reported in earlier studies (Zeng et al., 2018) (Figure S1C). This was also validated by immunostaining of X1, X2 and Xins cells for PIWI-1 protein, where X1 cells showed high PIWI-1 compared to X2 and Xins (Figures S2 D and E). Together these results show that the PIWI-1 antibody recognizes the neoblast population.

**Figure 2.**
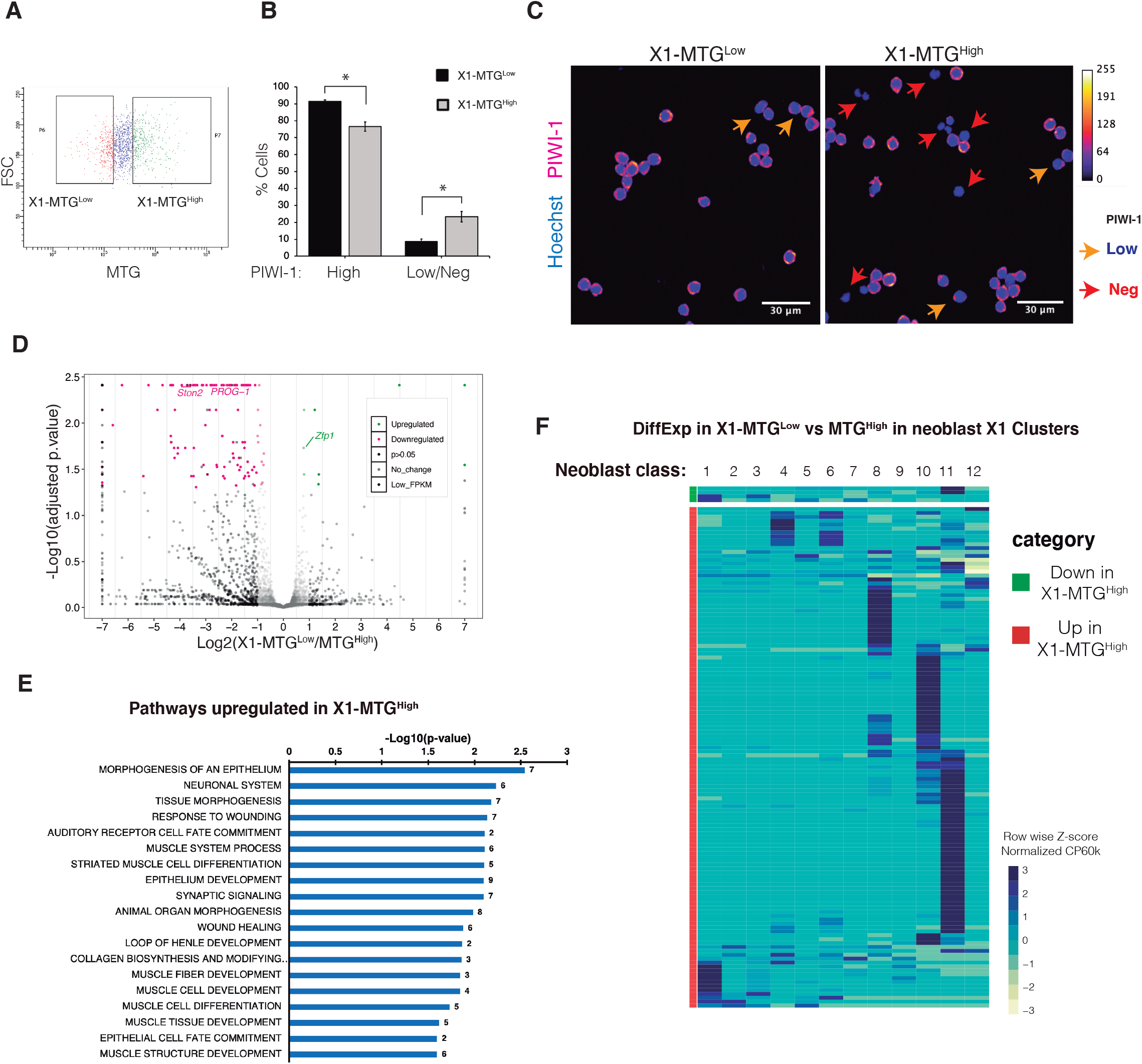
MTG staining reveals heterogeneity in X1 cells. **(A)** Dot plot showing the FACS gating for X1-MTG^Low^ and MTG^High^ cells. **(B)** PIWI-1 immunostaining in FACS sorted X1-MTG^Low^ and MTG^High^ cells quantified for PIWI-1^High^ through flow cytometry. Error bar represents s.e.m, N=3. * represents p<0.05. See also Figure S1. **(C)** Representative confocal images of PIWI-1 staining in indicated sorted population. PIWI-1 staining is shown using Fire LUT, (ImageJ). Arrow heads highlights PIWI-1 low (yellow) and negative (red) cells. Scale bar 30 μm. **(D)** Volcano plot displaying differential gene expression between X1-MTG^Low^ and MTG^High^ cells. **(E)** Bar graph representing selected GO terms that are downregulated in X1-MTG^Low^ cells compared to MTG^High^. Refer to supplemental file for full list of GO terms. See also Table S1. **(F)** Heatmap indicating the expression of the differentially expressed genes in X1-MTG^Low^ vs MTG^High^ cells in the published single cell database (Zeng et.al, Cell 2018).

Next, we examined the expression of PIWI-1 in MTG^High^ and MTG^Low^ cells within the X1 population (Figure 2A). We observed that X1-MTG^Low^ cells had higher representation of PIWI-1^High^ cells compared to X1-MTG^High^ cells (Figures 2B and C). Conversely, cells from X1-MTG^High^ cells were enriched in PIWI-1^Low/Negative^ cells. We speculate that an enrichment of PIWI-1^Low/Negative^ cells in X1-MTG^High^ could be the committed NB clusters and the X1-MTG^Low^ cells could either be the non-committed neoblast or the clonogenic population.

### Transcriptome sequencing of High and Low MTG reveal neoblast heterogeneity within X1 cells

In order to understand the functional states of High and Low MTG cells, RNA seq was performed on X1-MTG^Low^ and X1-MTG^High^ populations (Figure 2D). Transcriptome sequencing followed by Cuffdiff (Trapnell et al., 2013) analysis for differential expression identified 172 differentially regulated genes from X1-MTG^Low^ and MTG^High^ cells (Figure 2D). GO analysis of upregulated (p < 0.05) genes (163) in X1-MTG^High^ cells showed that most of these genes are essential for differentiation (muscle, epithelium, neural system), wound response and tissue morphogenesis (Figure 2E and Table S1). For instance, our analysis identified two key markers of epidermal and neural progenitors (*prog-1* and *ston-2)* upregulated by >2 fold (p < 0.05) in X1-MTG^High^ cells. Next, the differentially expressed genes were compared to the published single cell transcriptome data from X1 (Zeng et al., 2018). Most of the genes enriched in X1-MTG^High^ has high expression in NB classes other than NB2 which is the clonogenic neoblast population, suggesting that the X1-MTG^High^ cells are mostly committed neoblasts (Figure 2F). Interestingly, we did not observe any genes significantly upregulated in the X1-MTG^Low^ population that belong to the NB2 class (clonogenic neoblast) such as *tspan-1*, *tgs-1* and *pks-1*. Together, these results indicate that within X1, the MTG^High^ cells represent the committed neoblast and the MTG^Low^ cells could potentially contain pluripotent neoblast cells.

Encouraged by these results, we then explored the cell states of X2 population based on MTG signal. Cells with low and high MTG signal within X2 population were subjected to PIWI-1 immunostaining and transcriptome analysis.

### Low and High MTG X2 cells show heterogenous pool of PIWI-1^+^ and PIWI-1-populations respectively

Previous studies showed that X2 population consists of neoblasts and their progenitors (Eisenhoffer et al., 2008; Van Wolfswinkel et al., 2014). Our results based on MTG staining showed that X2 population can be broadly categorized as MTG^Low^ and MTG^High^ cells. To investigate the stem state of the MTG^Low^ and MTG^High^ cells in the X2 population, we stained with PIWI-1 antibody, which marks the neoblast populations. The X2-MTG^Low^ cells have mixed population of PIWI-1^High^ and ^-Low^. Flow cytometry analysis based on the size showed PIWI-1^High^ are large in size compared to the PIWI-1 low cells (Figure S1A) (Zeng et al., 2018). Hence, to separate PIWI-1^High^ cells in G1 phase, we further sorted the X2-MTG^Low^ and X2-MTG^High^ cells based on their size (FSC) which resulted in 4 populations (X2-MTG^Low^ High and Low FSC cells; X2-MTG^High^ High and Low FSC cells) (Figure 3A). The antibody staining indicated that within the X2-MTG^Low^, ~35% of HFSC (High FSC) cells showed higher expression of PIWI-1, while LFSC (Low FSC) cells had lower PIWI-1 expression (~85% cells) (Figure 3B and C). Regardless of the size, majority of the X2-MTG^High^ cells either showed low or negative expression for PIWI-1 (Figure 3B and C). Together, this data suggests that X2-MTG^Low^-(HFSC) cells, which were PIWI-1^High^, are likely to be neoblast population in G1 phase. Whereas the X2-MTG^Low^-(LFSC) cells, which are PIWI-1^Low^ could be the early progenitors. Conversely, X2-MTG^High^ (High and Low FSC) cells which are PIWI-1 negative could potentially be late progenitors.

**Figure 3.**
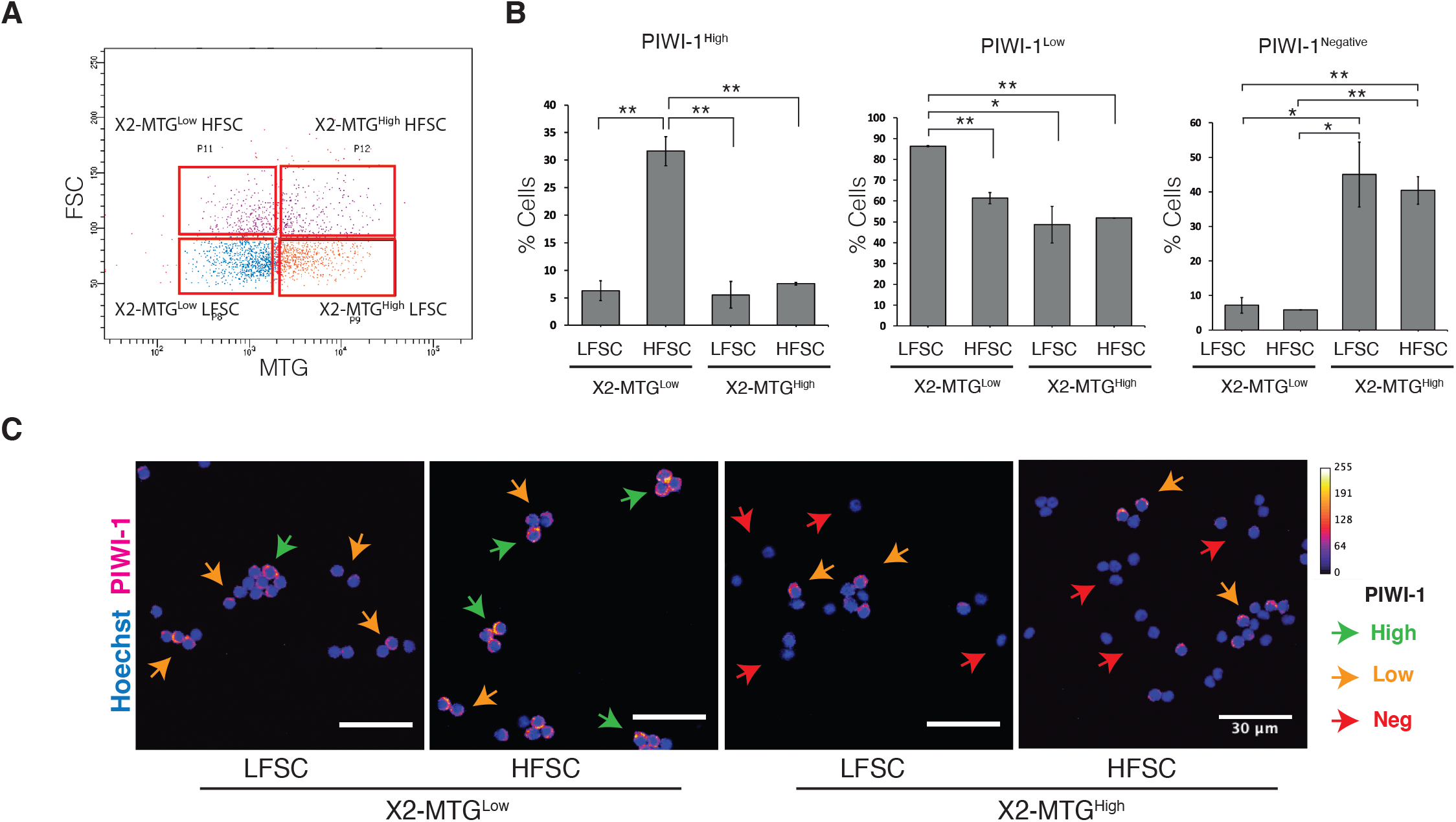
PIWI-1 immunostaining in X2-MTG subpopulations. **(A)** Dot plot representing the gating strategy for various MTG populations. HFSC - High Forward Scatter, LFSC – Low Forward Scatter. **(B)** Quantitative flow cytometry analysis of PIWI-1 staining in indicated FACS sorted X2 MTG population. N=3, *, ** represents p<0.05 and p<0.01 respectively. **(C)** Representative confocal images of PIWI-1 staining in indicated X2 MTG populations. PIWI-1 staining is represented in Fire LUT (ImageJ) and the pixel value and the corresponding color code is given in the graph (top right). Arrow heads indicates PIWI-1 high (green), low (yellow) and negative (red) cells. Scale bar: 30 μm. See also Figure S1

### Transcriptome sequencing show enrichment of neoblast and their early progenitors in X2-MTG^Low^ population

To gain further insights about the nature of the cells in the MTG^Low^ and MTG^High^ populations of X2, we performed RNAseq on the 4 populations. RNA seq analysis of X2-MTG^Low^ and MTG^High^ cells (High and Low FSC combined) revealed that the MTG^Low^ population is enriched with transcripts of stemness, nuage related neoblast markers (*Smedwi-1, Smedwi-3, Bruli*), cell cycle genes (*RRM2, MCM*) and early markers for differentiation compared to MTG^High^ (Figure 4A,B and Figure S2B,C and Table S2). Further, we also compared the RNA profile of the MTG subpopulations to the total X1, X2 and Xins populations isolated by conventional methods. Based on the principal component analysis, correlation plot and linkage mapping (Figures 4C and D, Figure S2A), the transcript signatures of X2-MTG^Low^ showed strong correlation (R^2^=0.8991) with X1 population compared to X2-MTG^High^, which was closer to Xins. Further, we also observed that most of the transcripts related to early epidermal progenitors (*prog-1* and *AGAT-1*) where enriched in X2-MTG^Low^, while the transcripts of late epidermal progenitors (*ZPUF6*) and the terminally differentiated epidermal cells were enriched in X2-MTG^High^ (Figures S2D). Moreover, pseudotime expression analysis using the available planarian single cell RNA seq database showed that the expression of transcripts enriched in X2-MTG^Low^ peaked in the neoblast class and decreased as the cells differentiated. In contrast, transcripts enriched in the X2-MTG^High^ showed negligible expression in neoblast populations and increased as the cells differentiate (Figures S3A-C). These analyses clearly show that the cells in X2-MTG^Low^ are either the neoblast cells in G1 phase or their immediate/early progenitors. Conversely, the cells in X2-MTG^High^ were mostly in the late phase of differentiation. Thus, our results suggest that the changes in mitochondrial content reflects the cell transition states within X2.

**Figure 4.**
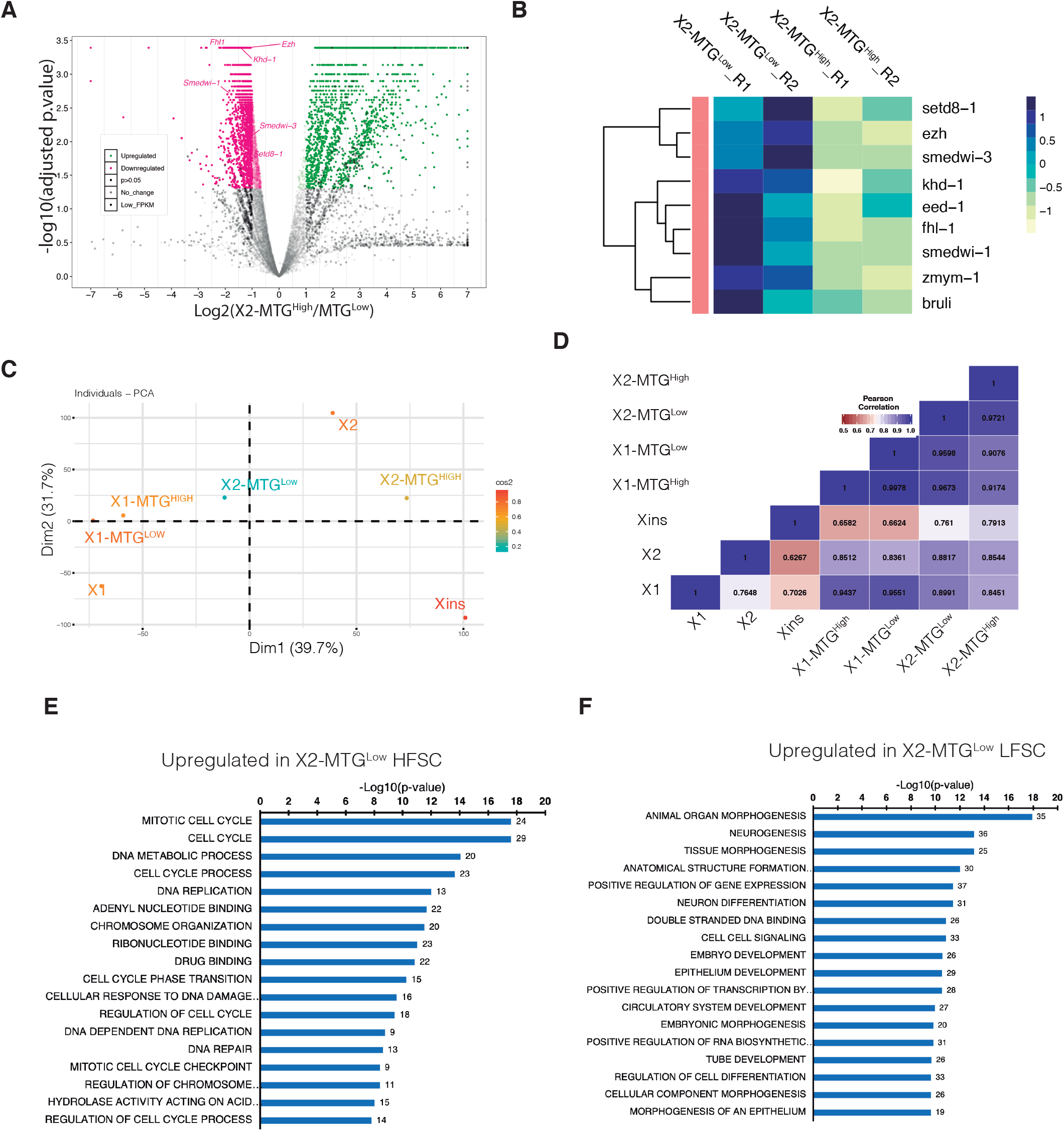
Transcriptome analysis of X2-MTG^Low^ and MTG^High^ cells. **(A)** Volcano plot showing the differentially expressed transcripts. Major stem cell genes >2 fold upregulated in MTG Low are marked in the plot. **(B)** Heat map showing expression of key stem cell related genes in X2-MTG^Low^ and MTG^High^ **(C)** Principal component analysis (PCA) showing indicated MTG population with respect to X1, X2 and Xins **(D)** Pearson’s correlation heatmap of the X2 MTG population compared with X1, X2 and Xins. See also Figure S2 and S3 and Table S2 **(E)** GO terms of statistically significant, >2fold upregulated transcripts from X2-MTG^Low^-HFSC cells compared to X2-MTG^Low^-LFSC cells. **(F)** GO terms of upregulated transcripts from the X2-MTG^Low^-LFSC cells compared to corresponding HFSC cells. See also Table S3

Next, we analyzed the transcriptome data from both High and Low FSC events from X2 MTG^Low^. GO analysis showed that the differentially expressed genes which are upregulated (p<0.05) in X2-MTG^Low^-(HFSC) cells are involved in cell cycle and DNA synthesis indicating these might be the stem cells in G0/G1 phase poised to potentially enter the cell cycle (Figure 4E). On the contrary, GO analysis for the transcripts enriched in X2-MTG^Low^-(LFSC) cells encode proteins that were involved in differentiation suggesting that these cells are the immediate/early progenitors (Figure 4F and Table S3). Further, the principle component analysis and correlation plot revealed that the transcripts from both the High and Low FSC cells of X2-MTG^Low^ show high correlation with X1 population (R^2^=0.9 HFSC cells and R^2^=0.89 LFSC cells) suggesting that X2-MTG^Low^-(HFSC) gate contains cells that are most likely the neoblast population in G1 phase (Figure S4A and B). Together, these results show that MTG along with Hoechst will be a good marker to identify and isolate neoblast cells in G0/G1 phase.

### Increased mitochondrial potential is essential for neoblast differentiation

To understand the mitochondrial activity of MTG^Low^ and ^-High^ cells, planarian cells were simultaneously stained with MTG and MitoTracker Orange (MTO), a dye which indicates mitochondrial membrane potential. It was observed that MTG^Low^ cells were also low in MTO and conversely MTG^High^ cells had high MTO intensity (Figure 5A and A’). Similar trend was also observed upon normalization of the MTO intensity to the cells treated with FCCP, an uncoupler of oxidative phosphorylation (Figure 5A and A’). This clearly indicate that the mitochondrial content correlates with mitochondrial activity. Next, we investigated the implication of changes in the mitochondrial activity on stem cell differentiation. To specifically perturb the mitochondrial activity in stem cells, we cultured neoblasts and treated with FCCP (Figure 5B). Towards this, we used SiR-DNA (Cytoskeleton inc.), a nuclear dye, recently used as an alternative to isolate 4N cells from planarians instead of Hoechst dye, which was shown to be toxic (Lei et al., 2019; Wagner et al., 2011; Wang et al., 2018b). The 4N cells which were also MTG^Low^ (X1-MTG^Low^ equivalent) was cultured as described by (Lei et al., 2019). The current cell culture system for neoblast is not completely optimized and it is observed that with time the cells differentiate and beyond 48 hours loose its clonogenic potential (Lei et al., 2019). We observed that post 48 hours of culture, there was higher number of PIWI-1^High^ cells in FCCP treated cells compared to the DMSO control (Figure 5C). This indicates that blocking OxPhos affects neoblast differentiation *in vitro*. Collectively, these results demonstrate that the differences in the mitochondrial content in the stem versus differentiated populations correlates with the mitochondrial activity and also suggests that an increased mitochondrial activity could be a requisite for neoblast differentiation.

**Figure 5.**
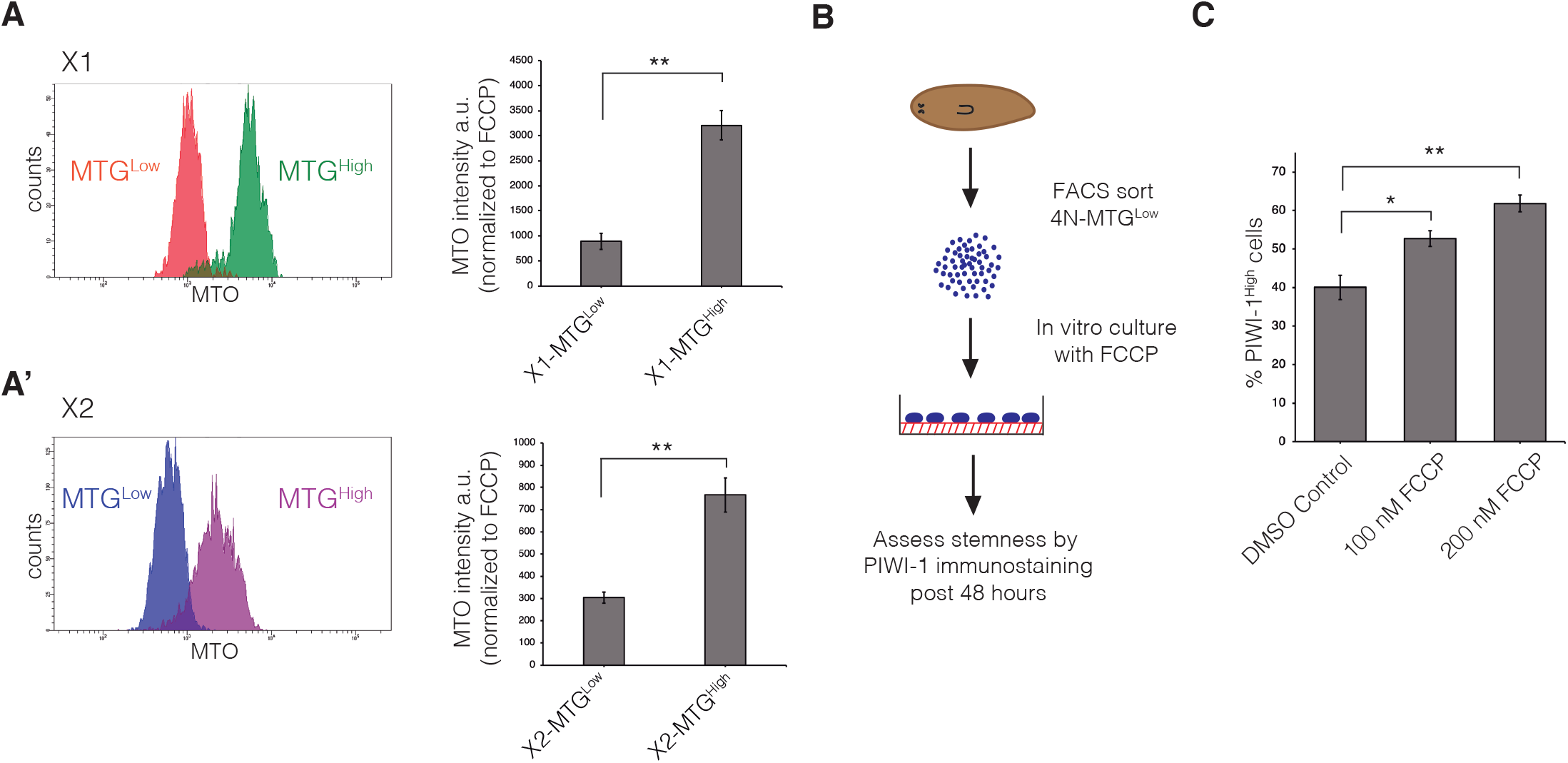
Mitochondrial potential measurement of X1 and X2 cells. **(A and A’)** Planarian cells stained with MTG and MTO analyzed through flow cytometry. X1 **(A)** and X2 **(A’)** were gated for MTG^Low^ and ^-High^ and analyzed for MTO intensity. Bar graph indicates median MTO intensity normalized to FCCP. N=3, error bar represents s.e.m. *, ** represents p<0.05 and p<0.01 respectively. a.u – arbitrary units **(B)** Schematic representing the *in vitro* assay for perturbing mitochondrial activity using FCCP **(C)** PIWI-1^High^ quantitation through flow cytometry in 4N-MTG^Low^ cells treated with indicated amount of FCCP for 48 hours in culture. N=3, error bar represents s.e.m. *, ** represents p<0.05 and p<0.01 respectively

### Pluripotency is associated with stem cells having low mitochondrial mass

The PIWI-1 expression and the transcriptome analysis revealed that the X1 and X2-MTG^Low^-(HFSC) cells could potentially be clonogenic neoblasts compared to their MTG^High^ counterparts. This was validated by transplantation experiment which involved injection of the MTG populations in to lethally irradiated animals. The population of cells that rescue the irradiated animals would be considered pluripotent stem cells (Lei et al., 2019; Wagner et al., 2011). Recent studies have shown that injection of neoblast from asexual into irradiated sexual planaria transformed sexual strain to asexuals (Lei et al., 2019; Wagner et al., 2011). For our transplantation experiments, we used SiR-DNA (Cytoskeleton Inc.) stained cells from asexual strain. First, the X1(FS) and X2(FS) gate was set using the Hoechst dye to isolate the 4N cells and 2N cells. The same gating parameters were used to separate the SiR-DNA 4N cells and SiR-DNA 2N cells to reduce the cross contamination between X1, X2 and Xins cells. The SiR-DNA cells were stained with MTG to demarcate the MTG^Low^ and MTG^High^ cells within the X1 and X2 population. To understand PSC activity of the MTG populations, we performed *piwi-1* colony expansion assay and animal survival studies wherein the 6 different populations of the sorted cells from asexual strain were transplanted into irradiated sexual strain. Animals that were not transplanted were used as a negative control. The animals after 8 days post transplantation (dpt) were fixed and assayed for the presence of *piwi-1*^+^ colonies (Figure 6A). We found that 4N-MTG^Low^ cells (X1-MTG^Low^ equivalent) had maximum number of *piwi-1*^*+*^ colonies compared to the 4N-MTG^High^ (Figure 6B). Similarly, among the 2N cells (X2 equivalent), only the 2N-MTG^Low^ (HFSC) cells showed greater number of *piwi-1*^*+*^ colony expansion (Figure 6B). None of the other three population (X2-MTG^Low^-LFSC, X2-MTG^High^ High and low FSC cells) showed any *piwi-1*^*+*^ colonies. We also performed survival studies in the transplanted and non-transplanted animals post lethal dose of irradiation. Typically, the non-transplanted sexual animals, post irradiation, show head regression, ventral curling and eventually lysed after 30 dpt. However, the sexual animals rescued with asexual neoblast eventually show fission, which is characteristic of asexual animals (Figures 6C and D). This result indicate that the transplanted cells have divided and replaced all the cells in the sexuals (Lei et al., 2019; Wagner et al., 2011). Within the 4N cells (X1 equivalent), MTG^Low^ population showed almost twice the rescue efficiency (65%) compared to 4N-MTG^High^ (35%) (Figure 6D and E). These results conclusively show that the neoblasts with low MTG are more pluripotent than the neoblasts with high MTG. Interestingly, the 2N-MTG^Low^-(HFSC) cells, showed 50% rescue efficiency compared to 2N-MTG^Low^ (LFSC) cells (5%, 1 out of 20 animals were rescued). As expected, both the 2N-MTG^High^ (small and large cells) failed to rescue the animals (Figure 6D and E). The new gating strategy described in this study, for the first time elucidates the presence of clonogenic neoblasts in X2 gate. Moreover, despite the presence of only ~35% PIWI-1^High^ cells, the X2-MTG^Low^-(HFSC) cells exhibited higher rescue efficiency compared to X1 MTG^High^ cells indicating a greater clonogenic potential (Figure S5A). Together our transplantation experiments show that the decreased mitochondrial mass as a hallmark of pluripotent stem cells in planarians. In contrast, the increased mitochondrial mass is an indicator of stem cell differentiation.

**Figure 6.**
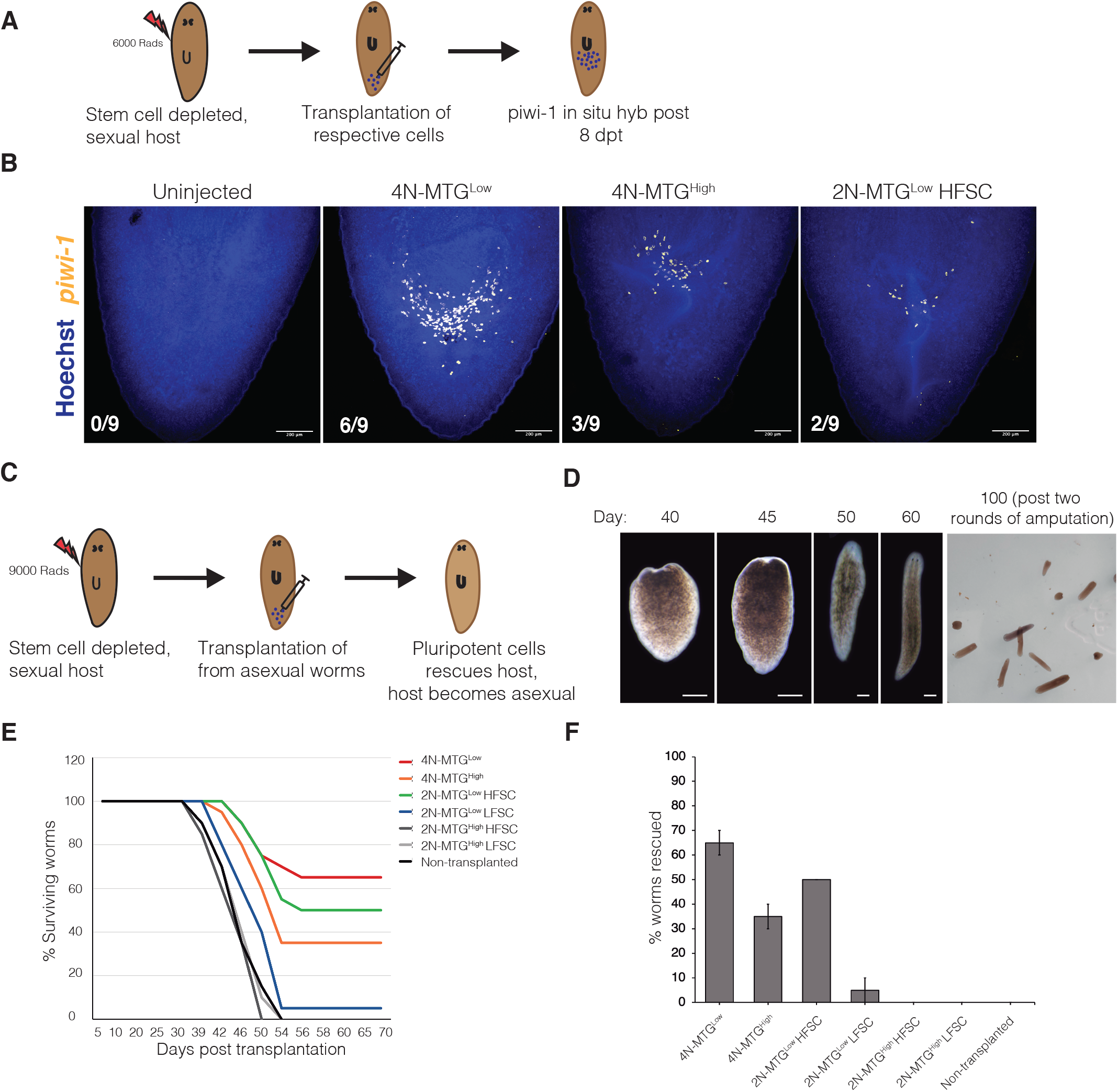
Transplantation experiment to validate the pluripotency of MTG populations. **(A)** Schematic of colony expansion assay performed **(B)** Representative *piwi-1* staining in lethally irradiated sexual worms 8 day post transplantation of ~1000 cells from asexual, SiR-DNA stained MTG populations (see methods). Numbers indicates worms showing *piwi-1* colonies. Scale bar: 200 μm. n=9 worms. **(C)** Schematic showing the long term survival and transformation of sexuals to asexuals by clonogenic neoblast transplantation. **(D)** Representative images of worms surviving after receiving clonogenic neoblasts. Initially the worms show head regression and around 40-45 dpt, worms start to develop blastema and completely regenerates head. After two rounds of amputation, the worms started propagating via fission. Scale bar, 500 μm. **(E)** Plots showing the percentage of surviving worms transplanted with ~1500 cells of the respective MTG population. n=20 worms. **(F)** Bar graph representing the average percentage of worms rescued. Error bar represents standard deviation, N=2, n=10 worms each. See also Figure S5

## Discussion

Planarians are highly regenerative animals that harbor pluripotent stem cells, which differentiate to lineage primed progenitors thereby controlling the extent of regeneration and tissue turnover during homeostasis (Reddien, 2018). The presence of pluripotent stem cells in the adult animal makes planaria an excellent model system to study stem cell dynamics. Earlier studies using electron microscopy had shown irregular mitochondrial morphology in the neoblast and most of the mitochondria were located towards the periphery of the nuclear membrane (Hay and Coward, 1975; Morita et al., 1969; Pedersen, 1959). Also, previous studies have shown an increase in the ROS (reactive oxygen species) activity during the planarian regeneration and it was critical for the differentiation of stem cells (Pirotte et al., 2015). This study was corroborated by the work in *Xenopus* and zebrafish, which showed increased ROS activity during their tail regeneration (Gauron et al., 2013; Love et al., 2013). Inhibiting the ROS activity using the NADH oxidase inhibitors abrogated the stem cell differentiation and thus blocked regeneration implicating mitochondria as the major source of ROS generation. The primary aim of this study was to delineate the changes in mitochondrial content in the pluripotent and the committed stem cell states in planarians, which can subsequently be used as a marker to isolate pluripotent stem cells.

Neoblast in planarians are characterized by the presence of large nucleus and scanty cytoplasm. Using Hoechst, a nuclear dye, planarian cells are sorted in to 3 classes, X1, X2 and Xins. Here, we investigated the extent of distribution of mitochondrial mass among these three populations using MTG, a mitochondrial dye. MTG accumulates in the mitochondrial matrix and covalently binds to the proteins by reacting with the free thiol groups of the cysteine residues. Its localization to the mitochondria happens regardless of the mitochondrial membrane potential and it has been used to measure the mitochondrial mass. MTG staining further demarcated X1 and X2 populations into MTG^Low^ and MTG^High^ subpopulations. We observed that PSCs in planaria are defined by low mitochondrial content whereas an increase in mitochondrial content is associated with differentiation. For instance, our results indicate that the stem cells in X1 population which had high mitochondrial content exhibited properties of lineage commitment such as reduced PIWI-1^High^ cells and expression of transcripts associated with lineage specification. This suggest that a quantifiable change in mitochondrial mass exists within clonogenic and specialized neoblasts.

We also looked at the mitochondrial content in X2 population, which is a mix of neoblasts and their mitotic progenies. Owing to the heterogeneity of the X2 population and low prevalence, stem cells in this gate were poorly studied. It was proposed that the clonogenic neoblasts exist in X2 gate but nonetheless it was never experimentally validated (Molinaro and Pearson, 2018). We categorized X2 cells based on MTG intensity and characterized the subpopulations. RNA seq analysis showed that the transcript profile of MTG^Low^ resembled stem cells and early progenitors and was reminiscent of X1 cells while MTG^High^ were more akin to Xins. This indicated that within X2, mitochondrial content increases as the cell progresses towards differentiation. To enrich the neoblasts within G0/G1 phase, MTG^Low^ cells in X2 were divided into two populations based on their size. Such classification of the X2 cells based on the size and MTG resulted in an enrichment of PIWI-1^High^ cells in High FSC gate. Transcriptome sequencing revealed that X2-MTG^Low^-(LFSC) represent early progenitors and the X2-MTG^Low^-(HFSC) express transcripts critical for the cell cycle progression. Together, using the size parameter and mitochondrial mass measurements, we were able to isolate neoblast and their immediate progenitors for functional characterization.

*Schmidtea mediterranea* exist as two strains; sexual strain, which are hermaphrodites and asexual strain, which has underdeveloped germline tissue and undergo reproduction by fission. Previous studies have shown that the transplantation of asexual neoblast to irradiated sexual strain transformed the sexual planaria to asexual evident from them undergoing fission (Wagner et al., 2011). Such transplantation assays in planarians provides a platform to functionally validate the pluripotency of neoblast subpopulations. Our results clearly indicates that stem cells with low mitochondrial content is more pluripotent than the cells with high mitochondrial content. This implies that the low mitochondrial content as one of the characteristics of pluripotent stem cells. Interestingly, our transplantation experiments also revealed higher clonogenic capacity of the G0/G1 neoblasts. With the current state of art technologies, TSPAN-1^+^ cells fairs as the most potent clonogenic population. We propose that classification of cells based on mitochondrial content would allow us to sort clonogenic neoblasts with higher purity and subsequent characterization might allow us to identify novel and rare clonogenic neoblasts, should they exist. Furthermore, the sorting methodology presented here using both nuclear and mitochondrial markers could be, in principle extended to other flatworms such as *Macrostomum lignano, Schistosoma mansoni* and *Hoefstenia miamia* where, like planaria, only proliferating stem cells have been extensively studied (Gehrke et al., 2019; Grudniewska et al., 2016; Wang et al., 2018a).

Differentiation in mammalian pluripotent cells such as ESCs are shown to involve changes in the gene expression profile, remodeling of organelles and their metabolic states (Dixon et al., 2015; Folmes et al., 2012; Gabut et al., 2020; Sampath et al., 2008). Further, it has been shown that during differentiation, ESCs exhibit increased mitochondrial fusion and a concurrent switch in their metabolic state from glycolytic to oxidative phosphorylation. Such metabolic change has been shown to be critical for the cell state transition (Chung et al., 2007; Tormos et al., 2011). For instance, a block in the electron transport chain using uncouplers like CCCP resulted in defective differentiation of ESCs (Mandal et al., 2011; Varum et al., 2009). Therefore, a change in the mitochondrial membrane potential was proposed as a marker for stemness vs differentiated states (Lonergan et al., 2007; Sukumar et al., 2016).

Our study demonstrates a substantial increase in the mitochondrial mass during differentiation and a perturbation of mitochondrial activity led to an increased sustenance of stem cells *in vitro*. This indicates that for proper differentiation of stem cells, a concomitant increase in the mitochondrial activity is a necessity. Further, these results suggest that the mitochondrial state has a central role in stem cell maintenance and differentiation in planarians and this biological process is evolutionarily conserved. Our findings also highlight the potential role of mitochondrial bioenergetics in regulating organismal regeneration. In addition, the increased mitochondrial content during differentiation might be a result of the higher demand for metabolites in cells priming for the differentiation. For instance, early stage embryonic stem cell differentiation is marked by epigenetic changes and increased translation state, which requires increased levels of amino acids and cofactors such as alpha-ketoglutarate and acetyl CoA (Boland et al., 2014; Folmes et al., 2012). Also, the intermediates of TCA cycle serve as the precursors of amino acid biosynthesis and also provide cofactors essential for epigenetic modifiers such as acetylases and demethylases. However, in depth study is required for complete understanding of the exact processes that leads to increased mitochondrial mass during neoblast differentiation. In summary, the results presented here establish without ambiguity that the pluripotent stem cells have lesser mitochondrial mass compared to their progenitors and this difference in the mitochondrial mass serves as an efficient marker to isolate pluripotent neoblast from their progenitors.

## Experimental Procedures

### Planarian husbandry

Both sexual and asexual *Schmidtea mediterranea* strains were grown in 1x Montjuïc salts at 20 degrees Celsius. The worms were fed with beef liver. The worms were starved for a minimum of 7 days before any experiments. Sexual strains receiving 6,000 rads of γ-rays were used as transplantation hosts. The transplantation hosts were maintained in gentamicin (50 μg/mL) starting 7 days prior to irradiation.

### Fluorescence-activated cell sorting

Cell suspension for FACS sorting was prepared as described before (Lei et al., 2019). The worms were diced in calcium and magnesium free buffer with 1% bovine serum albumin (CMFB) and mechanically sheared using a micropipette. The resulting single cell suspension was stained with Hoechst 33342 (40 μg/mL) and MTG (100 nM). X1, X2 and Xins population was demarcated using Hoechst blue and red fluorescence. For transplantation experiments, cells stained with SiR-DNA and MTG were used.

### Fluorescent in situ hybridization and immunostaining

In situ hybridization for assessing the *piwi-1* colonies in transplanted worms were performed as described earlier (King and Newmark, 2013). Digoxigenin labelled RNA probes were used for *piwi-1* detection. The signal was developed using Anti-digoxigenin POD antibody and CY-3 tyramide using tyramine signal amplification. For immunostaining, Anti-PIWI-1 antibody was raised in rabbit using the antigen NEPEGPTETDQSLS as described earlier (Guo et al., 2006). FACS sorted cells were plated in optical bottom 384 well plates (~10,000 cells per well) and stained with PIWI-1 antibody.

### Cell transplantation

Bulk cell transplantation in irradiated animals was carried out as described earlier with minor modifications (Davies et al., 2017; Wang et al., 2018b). For colony expansion and long term survival experiments, 2 days post irradiated animals were used. Irradiated worm was placed ventral side up above a black filter paper placed in a cold plate. The injection was carried out using an Eppendorf femtojet 4x with a pressure of 0.8-1.0 psi. For colony expansion assay, ~1000 cells/μL was injected and for long term survival assay ~1500 cells/μL was injected into the post gonopore midline of sexual hosts.

### Statistical analysis

Statistical significance of PIWI-1 antibody staining and MTO intensity was performed using unpaired, two tailed student’s TTEST. Flow cytometry analysis were performed in three independent biological replicates and P< 0.05 was considered to be statistically significant. Differentially expressed genes from the transcriptome analysis (two independent biological replicates) were identified using Cuffdiff module and genes with adjusted pvalue <0.05 were considered as significant. For GO analysis, >2 fold upregulated, statistically significant genes were considered.

## Data Availability

All the sequencing data generated in this study are uploaded in NCBI-Sequence Read Archive (SRA) under the accession id SRP272800.

## Acknowledgments

The authors thank Prof. Alejandro Sanchez Alvarado for hosting and training MMH on transplantation experiments in his laboratory at the Stower’s Institute for Medical Research. We thank Prof. Apurva Sarin for the valuable inputs and critical comments on the manuscript. We acknowledge all the Palakodeti lab members especially Srikar Krishna for the critical inputs. The authors thank Central Imaging and Flow Cytometry Facility (CIFF-BLiSc campus). MMH is supported by DBT JRF, VL is supported by CSIR-SRF, SRS thanks NCBS-PhD fellowship for graduate studies. The work was funded by DST Swarnajayanti Fellowship (DST/SJF/LSA-02/2015-16) awarded to DP and PKV acknowledges Department of Biotechnology, inStem for core funding.

## Author contributions

MMH, PKV and DP conceived and designed the study. MMH, SRS performed the experiments. VL performed the RNA seq analysis. KL, PKV and DP supervised the entire study. MMH and DP wrote the manuscript with inputs from all the authors. PKV and DP acquired funding.

## Supplemental Information

### Supplemental figure legends

**Supplementary Figure S1.**
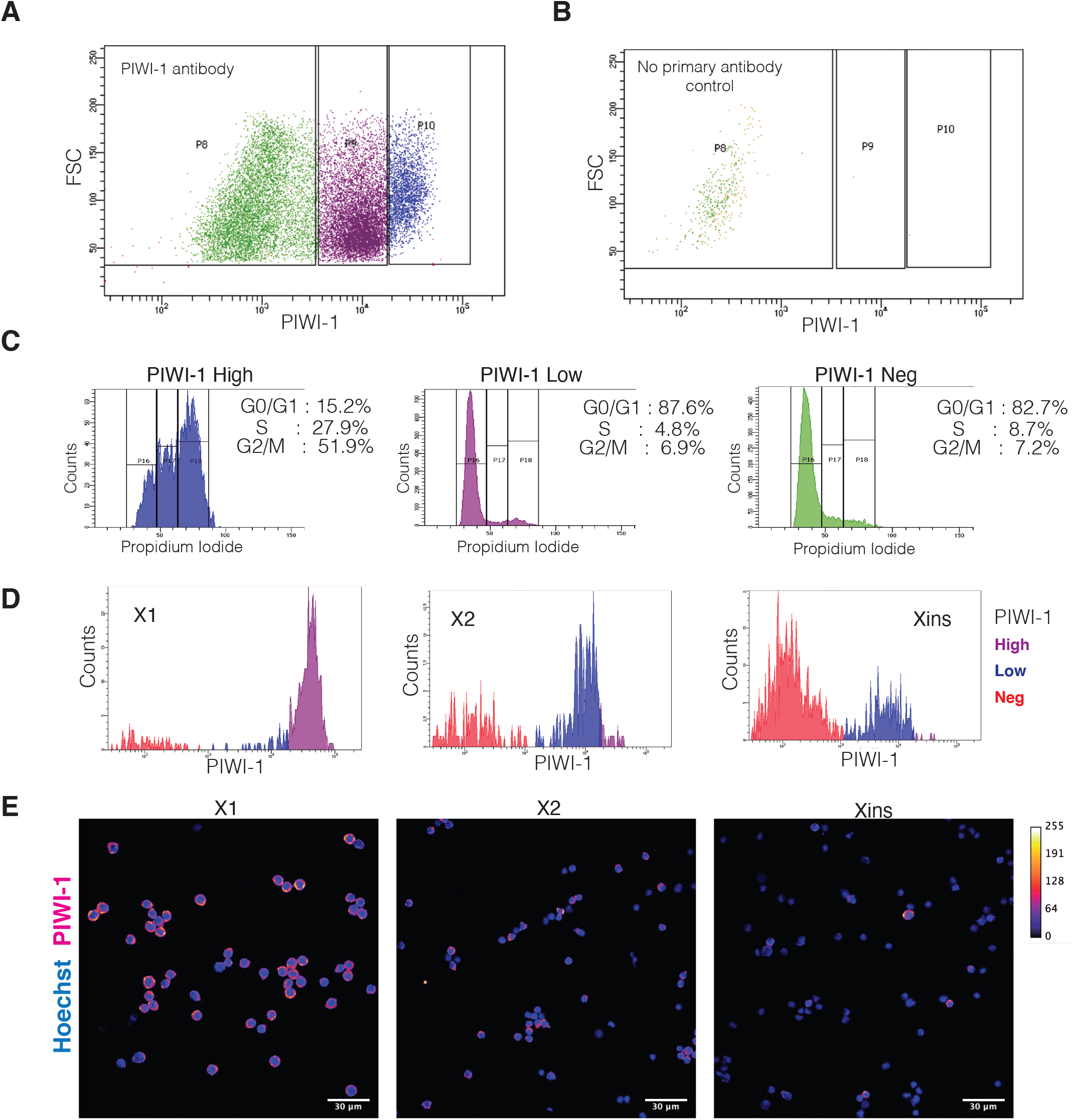
PIWI-1 Antibody staining validation (related to Figure 2,3) **(A)** Dot plot showing PIWI-1 staining in planarian cells. PIWI-1 High (blue), low (pink) and negative (green). **(B)** Dot plot of negative control cells-only secondary antibody was added without any primary antibody. **(C)** Cell cycle stages of corresponding PIWI-1 population analyzed by flow cytometry. Percentage of cells in G0/G1, S and G2/M for the indicated PIWI-1 population is given. **(D and E)** PIWI-1 staining in sorted X1, X2 and Xins cells analyzed by flow cytometry **(D)** or confocal microscopy **(E)** scale bar: 30 μm. PIWI-1 signal is shown as Fire LUT (ImageJ)

**Supplementary Figure S2.**
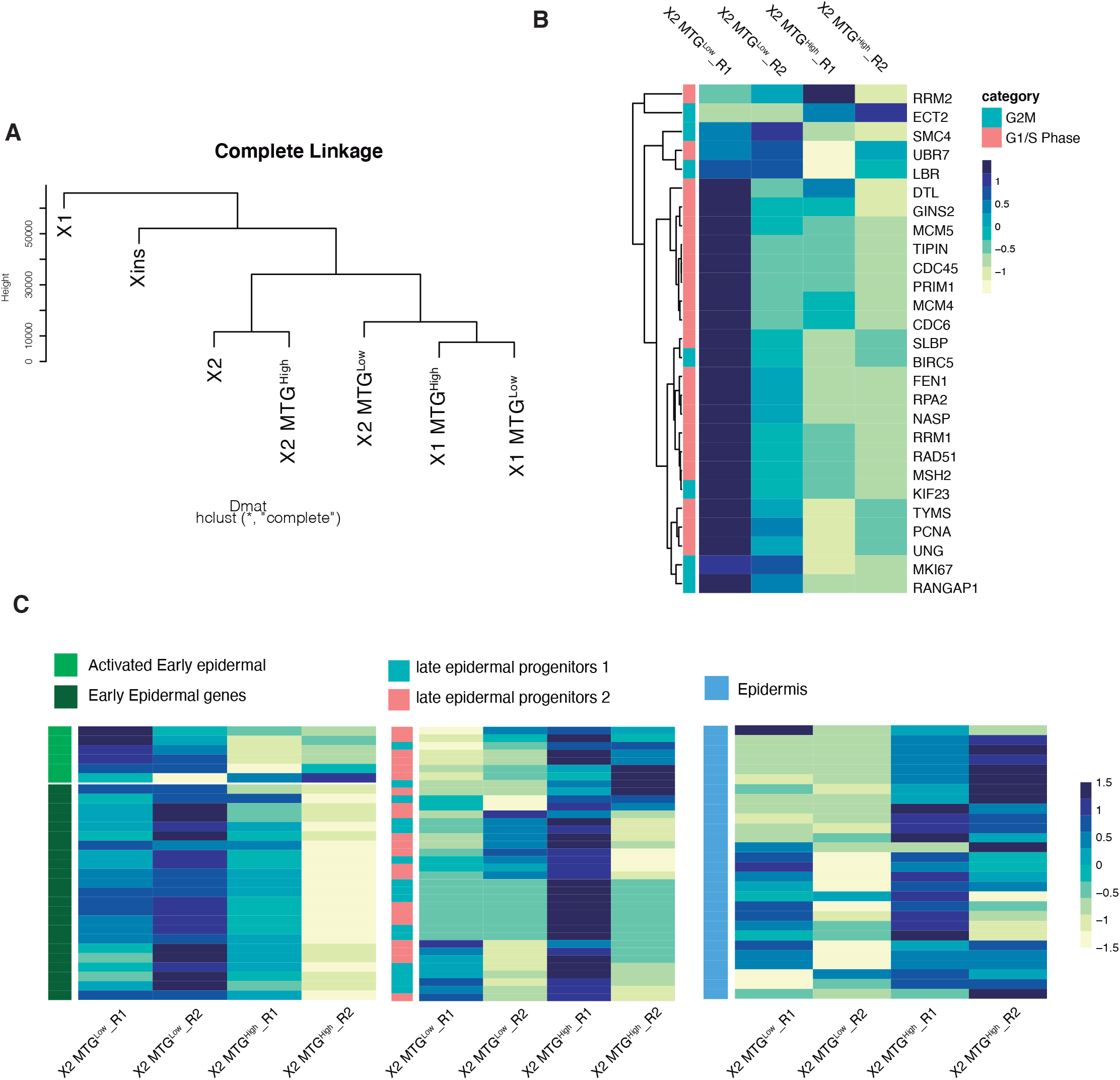
Transcriptome analysis of X2-MTG^Low^ and MTG^High^ population (related to Figure 4) **(A)** Linkage mapping of X2-MTG populations with X1-MTG and X1, X2 and Xins populations. **(B-C)** Heat map showing expression of key genes related to cell cycle (B) and (C) genes implicated in early epidermal progenitors, late progenitors and epidermis in X2-MTG^Low^ and MTG^High^ cells.

**Supplementary Figure S3.**
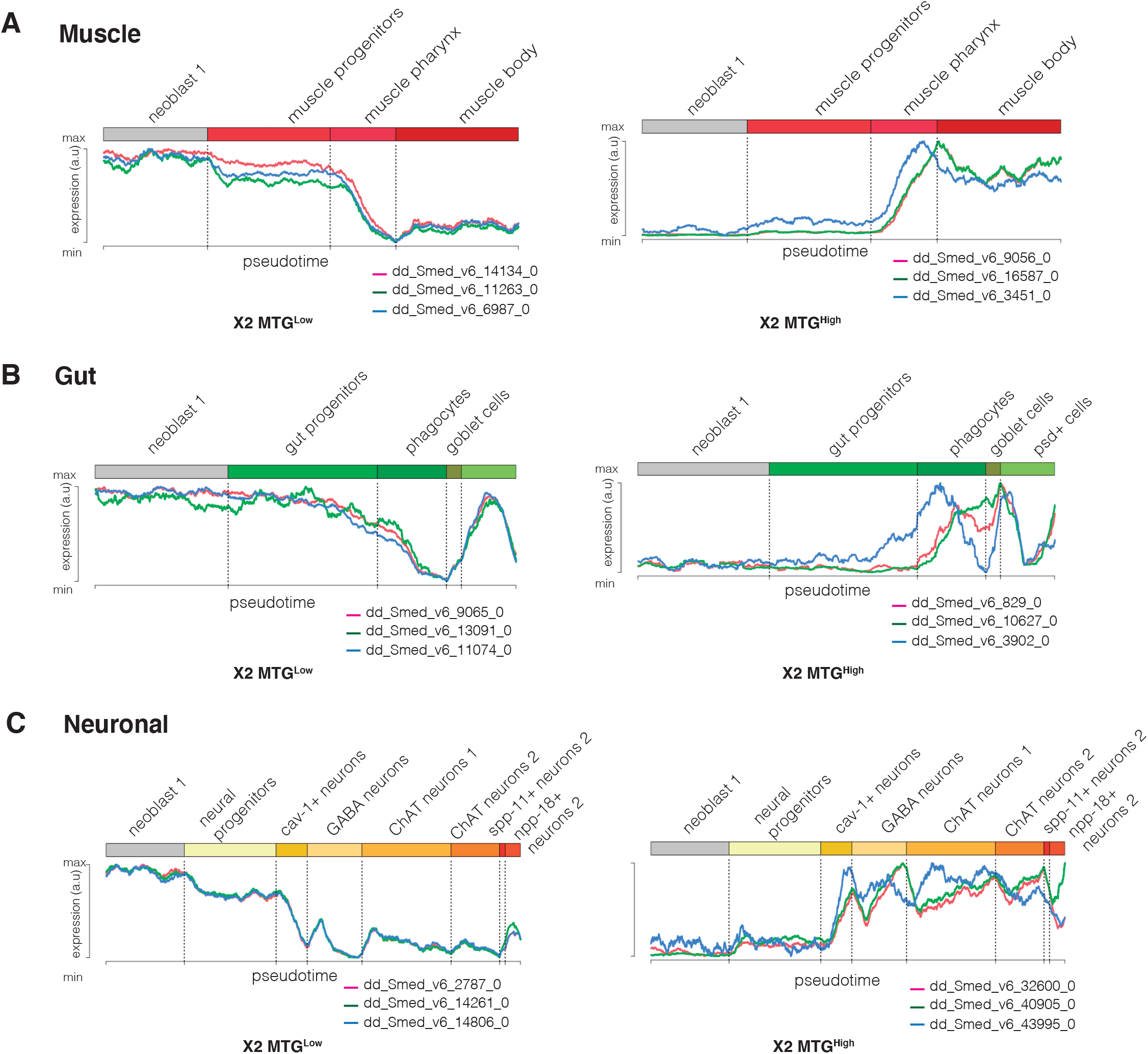
Pseudotime analysis of transcripts X2-MTG^Low^ and MTG^High^ population (related to Figure 4) **(A-C)** Representative transcripts enriched in either X2-MTG^Low^ (left) or X2-MTG^High^ (right) where used to analyze the pseudotime expression from neoblasts to terminally differentiated cells using available single cell database. Three representative transcripts related to muscle **(A)**, gut **(B)** and neuronal **(C)** lineages are shown

**Supplementary Figure S4.**
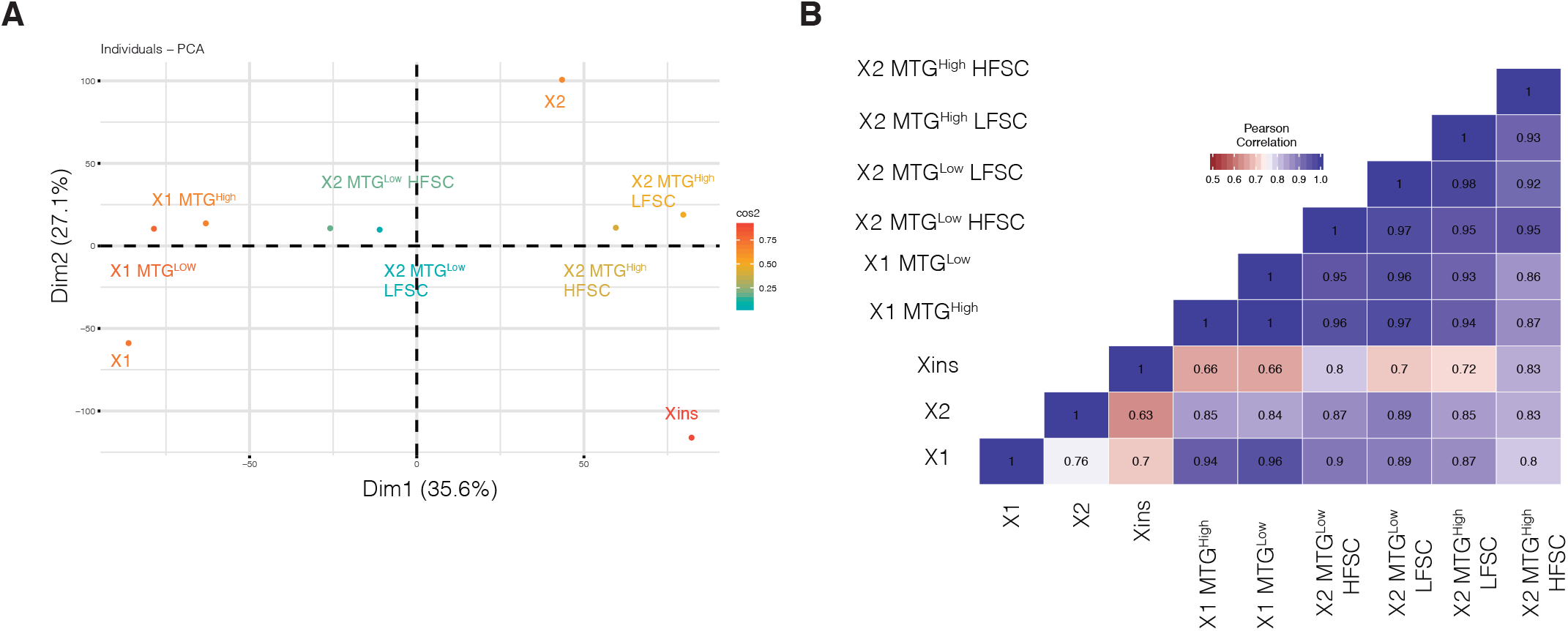
Transcriptome analysis of X2-MTG^Low^ High and Low FSC populations (related to Figure 4) **(A)** Principal component analysis comparing all the MTG subpopulations with X1, X2 and Xins **(B)** Pearson’s correlation heatmap of the MTG populations compared to X1, X2 and Xins

**Supplementary Figure S5.**
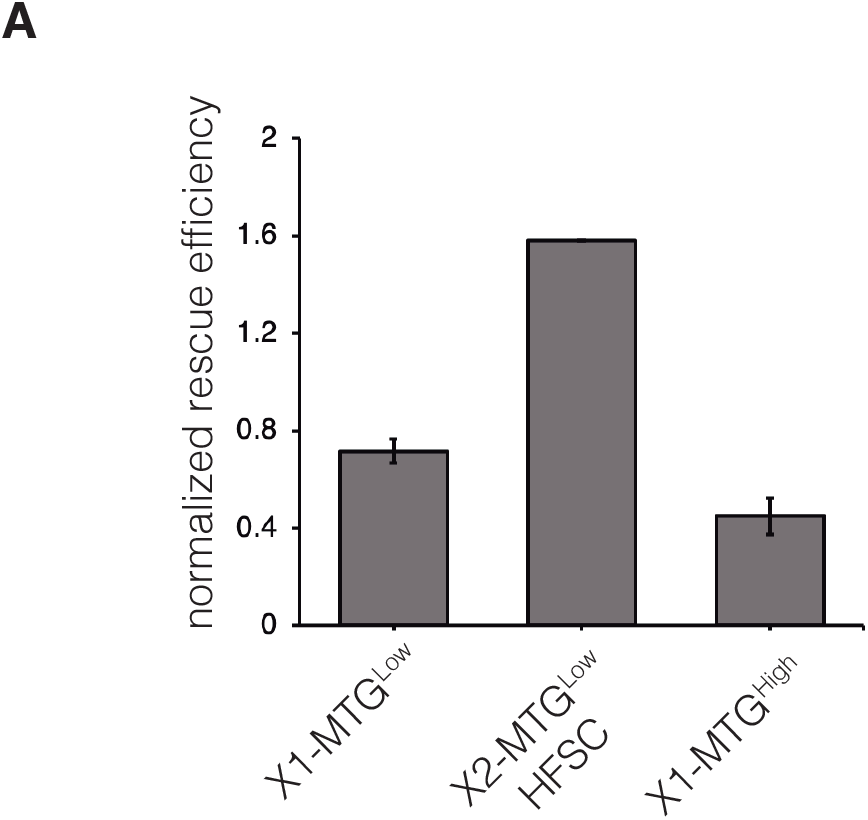
Normalized rescue of MTG subpopulations (related to Figure 6) **(A)** Normalized rescue efficiency calculated by dividing the rescue efficiency to the percentage of PIWI-1^High^ cells obtained from Figure 2B and 3B.

## Supplemental experimental procedures

### FACS sorting of cells using Hoechst 33342 and SiR-DNA

Cell suspension was prepared as described before (Lei et al., 2019). In brief, planarians were chopped in calcium and magnesium free buffer (Wang et al., 2018) with 1% bovine serum albumin (CMFB) (Sigma BSA A2153) and transferred to a 50 mL centrifuge tube using a wide bore pipette tip. The diced fragments are mechanically sheared by pipetting until all the fragments are dissociated. The resulting cell suspension is strained through 40 μm cell strainer (BD Falcon, 352340) and centrifuged at 290 g for 10 mins. Then, the supernatant was removed and isotonic planaria media (IPM) (Lei et al., 2019) with 10% FBS is added. To this solution, 40 μg/mL Hoechst 33342 (Invitrogen, H3570) was added and the staining is carried out in room temperature (RT) for 50 mins with intermittent mixing. To the same suspension, MitoTracker Green FM (Invitrogen, M7514) was added to a final concentration of 100 nM and stained for further 20 mins. Post staining, the suspension was again centrifuged (290 g, 10 mins) and the supernatant was discarded. The resulting cells were resuspended in IPM 10% FBS with 1 μg/mL propidium iodide (Sigma, P4864) and immediately analyzed through flow cytometry. Similar protocol was used for SiR-DNA (1 μM, Cytoskeleton, Inc. CY-SC007) staining, except for propidium iodide, DAPI (1 mg/mL) (Sigma D9542) was used to discriminate live/dead cells. For size based sorting of MTG populations, forward scatter voltage was adjusted such that X1 and X2 populations are well separated in FSC axis. First, Hoechst stained cells were used to define the forward and side scatter for X1(FS) and X2(FS) gates. Within the X1(FS) gate 4N SiR-DNA population was selected (X1 equivalent) and within X2(FS) gate 2N cells were selected (X2 equivalent) in order to reduce cross contamination. For Verapamil (Cytoskeleton, Inc. CY-SC007) treatment, the planarian cell suspension was treated with Hoechst and MTG in the presence of indicated concentrations of Verapamil for 1 hour. The cells were sorted using 100 μm tip and 20 psi sheath pressure in BD FACS ARIA III or Aria Fusion sorter.

### Quantitation of mitochondrial membrane potential

Planarian single cell suspensions prepared as described previously was treated with Hoechst for 40 mins and simultaneously stained with MTG and MTO (100 nM each) for another 20 mins and analyzed through flow cytometry (BD FACS Fortessa). As a control, cells were also treated with 20 μM of FCCP for 10 mins prior to addition of MTO. Median MTO intensity of FCCP treated cells were subtracted from the non-FCCP treated population to get the true membrane potential values.

### Antibody staining of FACS sorted cells

Anti-PIWI-1 antibody was raised in rabbit using the antigen NEPEGPTETDQSLS as described earlier (Guo et al., 2006). FACS sorted cells were centrifuged (300 g, 10 mins) and resuspended in IPM 5% FBS and 50 μL (~10,000 cells) was added in 384 well plates (Thermo Scientific Nunc, 142761). The plates were centrifuged and the cells were fixed in 50 μL, 4% PFA in PBSTx (0.1% Triton X100) for 10 minutes. Post incubation, 50 μL was removed and again 50 μL of fresh PFA fixative was added and incubated for further 10 mins. Then, the entire solution was carefully removed and washed with PBSTx and permeabilized with ice cold 90% methanol for 10 minutes. After washing twice with PBSTx, blocking was carried out for 30 minutes at RT with 10% horse serum in PBSTx. Anti-PIWI-1 antibody was diluted to 1:300 in blocking buffer was added and incubated overnight at 4 degree Celsius. After washing with PBSTx, secondary antibody was added (1:1000) (Anti rabbit IgG-Alexa Fluor 680, Invitrogen, A10043) for one hour at RT. The cells were further washed, and stained with Hoechst.

### Neoblast culture *in vitro*

Culturing of neoblast cells were performed as described previously (Lei et al., 2019). Briefly, ~120,000 FACS sorted 4N-MTG^Low^ cells were centrifuged and resuspended in KnockOut™ DMEM (Thermo Scientific 10829018) with 5% fetal bovine serum and plated in 24 well plate pretreated with Poly-D-Lysine (50 μg/mL, BD Biosciences, 354210). The media was supplemented with indicated concentration of FCCP or vehicle alone (DMSO) and cultured for 48 hours at 22 degree Celsius and 5% CO_2_. Post 48 hours, PIWI-1 antibody staining was performed as described earlier.

### Fluorescent in situ hybridization

In situ hybridization for assessing the *piwi-1* colonies in transplanted worms were performed as described earlier with minor modifications (King and Newmark, 2013). RNA riboprobe labelled with Digoxigenin against *piwi-1* was prepared by DIG RNA Labeling Mix (Sigma, 11277073910). 8 day post transplantation, worms were killed in 5% n-acetyl cysteine in PBS and fixed in 4% formaldehyde. The worms were serially dehydrated in methanol and stored in −20 degrees overnight. The worms were then rehydrated and bleached by H_2_O_2_ in formamide under white light. The bleached worms were permeabilized using Proteinase K (2 μg/mL), followed by fixation using 4% formaldehyde in PBSTx (0.5% Triton X-100). The worms were then transferred to prehybridization solution for 2 hour at 56 deg. Following this, hybridization was performed for 18-20 hours at 56 deg. Post hybridization, the worms were washed with 2X and 0.2X SSC buffer, followed by TNTx (0.1 M tris, 0.15 M NaCl, 0.3% Triton X-100, pH 7.5). The worms were then blocked in 5% horse serum, 0.5% Roche western blocking reagent in TNTx. Anti-digoxigenin POD antibody (1:1000) was added and incubated overnight at 4 degrees. The worms were then washed and developed using CY-3 tyramide signal amplification. The worms were counterstained with Hoechst and mounted using Scale A2.

### Microscopy

Olympus FV3000 and FV 1000 confocal microscopes were used to image *piwi-1* FISH and immunostaining. The images were analyzed using imageJ (https://imagej.nih.gov/ij). Micrographs of live worms recovering from irradiation experiments were acquired using Olympus SZ-16 stereo microscope.

### RNA extraction

Total RNA was extracted from sorted cell using Trizol™ method (Invitrogen, 15596026). Cells were centrifuged and 0.5 mL of Trizol™ was added for ~100,000 cells. RNA extraction was carried out as per manufacturer’s protocol. Total RNA extracted was quantified using Qubit RNA HS assay kit (Invitrogen Q32852) method and the RNA quality and integrity was verified using Bioanalyzer.

### Transcriptomic profiling of mitochondrial populations

Transcriptome profiling for 6 population (X1-MTG^Low^, X1-MTG^High^, X2-MTG^Low^ HFSC and LFSC cells, X2-MTG^High^ HFSC and LFSC cells) were performed. Transcriptome library was prepared using NEBNext® Ultra™ II Directional RNA Library Prep with Sample Purification Beads (Catalog no-E7765L) kit and sequenced in Illumina HiSeq 2500 machine. All the samples were sequenced (as single-end) in biological replicates. Approximately 25 to 40 million reads were sequenced for every sample. These reads were adapter trimmed using Trimmomatic (Bolger et al., 2014) and mapped to rRNA and other contamination databases. Reads that did not align to these databases were taken for further analysis. We used reference based transcriptome assembly algorithms Hisat2 v2.1.0 (Kim et al., 2015); Cufflinks v2.2.1 (Trapnell et al., 2010) and Cuffdiff v2.2.1 (Trapnell et al., 2013) to identify differentially expressed transcripts. We used Hisat2 (-q -p 8 --min-intronlen 50 --max-intronlen 250000 --dta-cufflinks --new-summary --summary-file) to align the reads back to dd_Smes_G4 (Grohme et al., 2018) assembly of *Schmidtea mediterranea* genome. Around 52-70% of reads were mapped to dd_Smes_G4 genome. We used samtools to obtain sorted bam files. The mapped reads were assembled using Cufflinks (-p -o -b -u -N --total-hits-norm -G) with most recent and well annotated SMEST transcriptome as reference (http://planmine.mpi-cbg.de/planmine/assemblyReport.do?assemblyId=SMEST.1_report.html). We used cuffmerge to merge the gene list across different conditions. We identified differentially expressed genes using Cuffdiff module (-p -o ./ -b -u -N --total-hits-norm -L) and considered genes with adjusted pvalue <0.05 as significance cut-off. Genes with significant p-value and atleast two fold up/down regulation were considered for GO and pathway analysis. Along with these mitochondrial based sub-populations we have used RNASeq data for total X1, X2 and Xins population from SRA. We downloaded X1 and Xins transcriptome data from <u>PRJNA296017</u> (four replicates each) (Tu et al., 2015) and X2 from <u>PRJNA167022</u> (two replicates) (Labbé et al., 2012). We analysed these data as described above and used for dimensional reduction analysis (PCA, clustering). Like X1 cells, to obtain only MTG low and high populations for X2, we considered HFSC and LFSC cells of same MTG levels (either low of high) as replicates. One of the replicate for X2 MTG High HFSC replicate showed poor correlation to genome, so we have removed that one replicate from the analysis.

We did pathway analysis & gene-ontology analysis for these selected up/down regulated transcripts using GSEA (Mootha et al., 2003; Subramanian et al., 2005). We used customized perl script for all the analysis used in this study. We used R ggplot2 (Wickham, 2016), pHeatmap and CummeRbund (Goff et al., 2012) library for plotting. Different planarian cell-type markers were obtained from the available single cell transcriptome data (Fincher et al., 2018; Plass et al., 2018; Van Wolfswinkel et al., 2014). Expression of these markers (FPKM values) from all the different cell populations are plotted as heatmaps using pHeatmap (https://cran.r-project.org/web/packages/pheatmap/index.html). Pseudotime analysis was performed with representative transcripts from different population using available single cell dataset (https://shiny.mdc-berlin.de/psca/)

### Bulk cell transplantation

Bulk cell transplantation in irradiated animals was carried out as described earlier with minor modifications (Davies et al., 2017; Wang et al., 2018). FACS sorted cells were centrifuged and resuspended in IPM 5% FBS media and kept in ice throughout the experiments. For colony expansion and long term survival experiments, 2 days post irradiated animals were used. Irradiated worm was placed ventral side up above a black filter paper placed in a cold plate. The injection was carried out using an Eppendorf femtojet 4x with a pressure of 0.8-1.0 psi. Glass capillaries (length 3.5”, ID: 0.53 mm, OD: 1.14 mm, Drummond Scientific, Inc. USA) were pulled using Sutter Instrument model P-1000. Cell suspensions were loaded onto the pulled capillaries using a mouth pipette (Sigma, A5177-5EA). For colony expansion assay, ~1000 cells/μL were injected and for long term survival assay ~1500 cells/μL was injected into the post gonopore midline of sexual hosts. The transplant hosts and the uninjected control worms was maintained in gentamicin 50 μg/mL in 6 well plates with planaria water changes every two days. After 60 days, the surviving worms were fed with beef liver and amputated to regenerate twice. Fission activity of the rescued worms were monitored.

## References

Boland, M.J., Nazor, K.L., and Loring, J.F. (2014). Epigenetic regulation of pluripotency and differentiation. Circ. Res.

Chung, S., Dzeja, P.P., Faustino, R.S., Perez-Terzic, C., Behfar, A., and Terzic, A. (2007). Mitochondrial oxidative metabolism is required for the cardiac differentiation of stem cells. Nat. Clin. Pract. Cardiovasc. Med.

Davies, E.L., Lei, K., Seidel, C.W., Kroesen, A.E., Mckinney, S.A., Guo, L., Robb, S.M.C., Ross, E.J., Gotting, K., and Sa, A. (2017). Embryonic origin of adult stem cells required for tissue homeostasis and regeneration. 1–35.

Dixon, J.R., Jung, I., Selvaraj, S., Shen, Y., Antosiewicz-Bourget, J.E., Lee, A.Y., Ye, Z., Kim, A., Rajagopal, N., Xie, W., et al. (2015). Chromatin architecture reorganization during stem cell differentiation. Nature.

Doherty, E., and Perl, A. (2017). Measurement of Mitochondrial Mass by Flow Cytometry during Oxidative Stress. React. Oxyg. Species.

Eisenhoffer, G.T., Kang, H., and Alvarado, A.S. (2008). Molecular Analysis of Stem Cells and Their Descendants during Cell Turnover and Regeneration in the Planarian Schmidtea mediterranea. Cell Stem Cell 3, 327–339.

Fincher, C.T., Wurtzel, O., de Hoog, T., Kravarik, K.M., and Reddien, P.W. (2018). Cell type transcriptome atlas for the planarian Schmidtea mediterranea. Science (80-.). 360.

Folmes, C.D.L., Dzeja, P.P., Nelson, T.J., and Terzic, A. (2012). Metabolic plasticity in stem cell homeostasis and differentiation. Cell Stem Cell.

Gabut, M., Bourdelais, F., and Durand, S. (2020). Ribosome and Translational Control in Stem Cells. Cells.

Gauron, C., Rampon, C., Bouzaffour, M., Ipendey, E., Teillon, J., Volovitch, M., and Vriz, S. (2013). Sustained production of ROS triggers compensatory proliferation and is required for regeneration to proceed. Sci. Rep.

Gehrke, A.R., Neverett, E., Luo, Y.J., Brandt, A., Ricci, L., Hulett, R.E., Gompers, A., Graham Ruby, J., Rokhsar, D.S., Reddien, P.W., et al. (2019). Acoel genome reveals the regulatory landscape of whole-body regeneration. Science (80-.).

Grudniewska, M., Mouton, S., Simanov, D., Beltman, F., Grelling, M., De Mulder, K., Arindrarto, W., Weissert, P.M., van der Elst, S., and Berezikov, E. (2016). Transcriptional signatures of somatic neoblasts and germline cells in Macrostomum lignano. Elife.

Guo, T., Peters, A.H.F.M., and Newmark, P.A. (2006). A bruno-like Gene Is Required for Stem Cell Maintenance in Planarians. Dev. Cell.

Hay, E.D., and Coward, S.J. (1975). Fine structure studies on the planarian, Dugesia. I. Nature of the “neoblast” and other cell types in noninjured worms. J. Ultrasructure Res.

Hayashi, T., Asami, M., Higuchi, S., Shibata, N., and Agata, K. (2006). Isolation of planarian X-ray-sensitive stem cells by fluorescence-activated cell sorting. 371–380.

King, R.S., and Newmark, P.A. (2013). In situ hybridization protocol for enhanced detection of gene expression in the planarian Schmidtea mediterranea. BMC Dev. Biol.

Lei, K., McKinney, S.A., Ross, E.J., Lee, H.-C., and Alvarado, A.S. (2019). Cultured pluripotent planarian stem cells retain potency and express proteins from exogenously introduced mRNAs. BioRxiv.

Lonergan, T., Bavister, B., and Brenner, C. (2007). Mitochondria in stem cells. Mitochondrion.

Love, N.R., Chen, Y., Ishibashi, S., Kritsiligkou, P., Lea, R., Koh, Y., Gallop, J.L., Dorey, K., and Amaya, E. (2013). Amputation-induced reactive oxygen species are required for successful Xenopus tadpole tail regeneration. Nat. Cell Biol.

Mandal, S., Lindgren, A.G., Srivastava, A.S., Clark, A.T., and Banerjee, U. (2011). Mitochondrial function controls proliferation and early differentiation potential of embryonic stem cells. Stem Cells 29, 486–495.

Molinaro, A.M., and Pearson, B.J. (2016). In silico lineage tracing through single cell transcriptomics identifies a neural stem cell population in planarians. Genome Biol.

Molinaro, A.M., and Pearson, B.J. (2018). Myths vs. FACS: What do we know about planarian stem cell lineages? Int. J. Dev. Biol.

Morita, M., Best, J.B., and Noel, J. (1969). Electron microscopic studies of planarian regeneration. I. Fine structure of neoblasts in Dugesia dorotocephala. J. Ultrasructure Res.

Newmark, P.A., and Sánchez Alvarado, A. (2000). Bromodeoxyuridine specifically labels the regenerative stem cells of planarians. Dev. Biol.

Pedersen, K.J. (1959). Cytological studies on the planarian neoblast. Zeitschrift Für Zellforsch. Und Mikroskopische Anat.

Pirotte, N., Stevens, A.S., Fraguas, S., Plusquin, M., Van Roten, A., Van Belleghem, F., Paesen, R., Ameloot, M., Cebrià, F., Artois, T., et al. (2015). Reactive oxygen species in planarian regeneration: An upstream necessity for correct patterning and brain formation. Oxid. Med. Cell. Longev.

Plass, M., Solana, J., Alexander Wolf, F., Ayoub, S., Misios, A., Glažar, P., Obermayer, B., Theis, F.J., Kocks, C., and Rajewsky, N. (2018). Cell type atlas and lineage tree of a whole complex animal by single-cell transcriptomics. Science (80-.). 360.

Reddien, P.W. (2018). The Cellular and Molecular Basis for Planarian Regeneration. Cell.

Reddien, P.W., and Alvarado, A.S. (2004). FUNDAMENTALS OF PLANARIAN REGENERATION. Annu. Rev. Cell Dev. Biol.

Reddien, P.W., Oviedo, N.J., Jennings, J.R., Jenkin, J.C., and Sánchez Alvarado, A. (2005). Developmental biology: SMEDWI-2 is a PIWI-like protein that regulates planarian stem cells. Science (80-.).

Romero-Moya, D., Bueno, C., Montes, R., Navarro-Montero, O., Iborra, F.J., López, L.C., Martin, M., and Menendez, P. (2013). Cord blood-derived CD34+ hematopoietic cells with low mitochondrial mass are enriched in hematopoietic repopulating stem cell function. Haematologica 98, 1022–1029.

Sampath, P., Pritchard, D.K., Pabon, L., Reinecke, H., Schwartz, S.M., Morris, D.R., and Murry, C.E. (2008). A Hierarchical Network Controls Protein Translation during Murine Embryonic Stem Cell Self-Renewal and Differentiation. Cell Stem Cell.

Sukumar, M., Liu, J., Mehta, G.U., Patel, S.J., Roychoudhuri, R., Crompton, J.G., Klebanoff, C.A., Ji, Y., Li, P., Yu, Z., et al. (2016). Mitochondrial Membrane Potential Identifies Cells with Enhanced Stemness for Cellular Therapy. Cell Metab. 23, 63–76.

Tormos, K. V., Anso, E., Hamanaka, R.B., Eisenbart, J., Joseph, J., Kalyanaraman, B., and Chandel, N.S. (2011). Mitochondrial complex III ROS regulate adipocyte differentiation. Cell Metab.

Trapnell, C., Hendrickson, D.G., Sauvageau, M., Goff, L., Rinn, J.L., and Pachter, L. (2013). Differential analysis of gene regulation at transcript resolution with RNA-seq. Nat. Biotechnol.

Varum, S., Momčilović, O., Castro, C., Ben-Yehudah, A., Ramalho-Santos, J., and Navara, C.S. (2009). Enhancement of human embryonic stem cell pluripotency through inhibition of the mitochondrial respiratory chain. Stem Cell Res.

Wagner, D.E., Wang, I.E., and Reddien, P.W. (2011). Clonogenic neoblasts are pluripotent adult stem cells that underlie planarian regeneration. Science (80-.).

Wagner, D.E., Ho, J.J., and Reddien, P.W. (2012). Genetic regulators of a pluripotent adult stem cell system in planarians identified by RNAi and clonal analysis. Cell Stem Cell 10, 299–311.

Wang, B., Lee, J., Li, P., Saberi, A., Yang, H., Liu, C., Zhao, M., and Newmark, P.A. (2018a). Stem cell heterogeneity drives the parasitic life cycle of Schistosoma Mansoni. Elife.

Wang, I.E., Wagner, D.E., and Reddien, P.W. (2018b). Clonal analysis of planarian stem cells by subtotal irradiation and single-cell transplantation. In Methods in Molecular Biology, p.

Van Wolfswinkel, J.C., Wagner, D.E., and Reddien, P.W. (2014). Single-cell analysis reveals functionally distinct classes within the planarian stem cell compartment. Cell Stem Cell 15, 326–339.

Xu, X., Duan, S., Yi, F., Ocampo, A., Liu, G.H., and Izpisua Belmonte, J.C. (2013). Mitochondrial regulation in pluripotent stem cells. Cell Metab. 18, 325–332.

Zeng, A., Li, H., Guo, L., Gao, X., McKinney, S., Wang, Y., Yu, Z., Park, J., Semerad, C., Ross, E., et al. (2018). Prospectively Isolated Tetraspanin+Neoblasts Are Adult Pluripotent Stem Cells Underlying Planaria Regeneration. Cell 1593–1608.

## References

Bolger, A.M., Lohse, M., and Usadel, B. (2014). Trimmomatic: A flexible trimmer for Illumina sequence data. Bioinformatics.

Goff, L.A., Trapnell, C., and Kelley, D. (2012). CummeRbund: visualization and exploration of Cufflinks high-throughput sequencing data. R Packag. Version.

Grohme, M.A., Schloissnig, S., Rozanski, A., Pippel, M., Young, G.R., and Winkler, S. (2018). The genome of Schmidtea mediterranea and the evolution of core cellular mechanisms. Nat. Publ. Gr.

Kim, D., Langmead, B., and Salzberg, S.L. (2015). HISAT: A fast spliced aligner with low memory requirements. Nat. Methods.

Labbé, R.M., Irimia, M., Currie, K.W., Lin, A., Zhu, S.J., Brown, D.D.R., Ross, E.J., Voisin, V., Bader, G.D., Blencowe, B.J., et al. (2012). A Comparative transcriptomic analysis reveals conserved features of stem cell pluripotency in planarians and mammals. Stem Cells.

Mootha, V.K., Lindgren, C.M., Eriksson, K.F., Subramanian, A., Sihag, S., Lehar, J., Puigserver, P., Carlsson, E., Ridderstråle, M., Laurila, E., et al. (2003). PGC-1α-responsive genes involved in oxidative phosphorylation are coordinately downregulated in human diabetes. Nat. Genet.

Subramanian, A., Tamayo, P., Mootha, V.K., Mukherjee, S., Ebert, B.L., Gillette, M.A., Paulovich, A., Pomeroy, S.L., Golub, T.R., Lander, E.S., et al. (2005). Gene set enrichment analysis: A knowledge-based approach for interpreting genome-wide expression profiles. Proc. Natl. Acad. Sci. U. S. A.

Trapnell, C., Williams, B.A., Pertea, G., Mortazavi, A., Kwan, G., Van Baren, M.J., Salzberg, S.L., Wold, B.J., and Pachter, L. (2010). Transcript assembly and quantification by RNA-Seq reveals unannotated transcripts and isoform switching during cell differentiation. Nat. Biotechnol.

Tu, K.C., Cheng, L.C., Vu, H.T.K., Lange, J.J., McKinney, S.A., Seidel, C.W., and Sánchez Alvarado, A. (2015). Egr-5 is a post-mitotic regulator of planarian epidermal differentiation. Elife.

Wang, I.E., Wagner, D.E., and Reddien, P.W. (2018). Clonal analysis of planarian stem cells by subtotal irradiation and single-cell transplantation. In Methods in Molecular Biology, p.

Wickham, H. (2016). ggplot2 Elegant Graphics for Data Analysis (Use R!).

